# Daikenchuto modulates gut microbial metabolism to mitigate irinotecan-induced enterotoxicity

**DOI:** 10.64898/2026.05.22.727323

**Authors:** Yinyue Xu, Rui Liang, Peng Xia, Siqian Luo, Bingyue Jiang, Anxin Wang, Kangli Liang, Yitao Wang, Wanghui Jing, Sicen Wang

## Abstract

Gut microbiota metabolic remodeling is a pivotal determinant in irinotecan-induced enterotoxicity and epithelial damage, although the underlying mechanisms remain unclear. Herein, we discovered that Daikenchuto (DKT), a traditional Chinese prescription for intestinal disorders, alleviated irinotecan–induced enterotoxicity without compromising its anti-tumor efficacy by improving weight loss, diarrhea, intestinal inflammation, and barrier damage, and these effects were partially dependent on gut microbiota. DKT significantly restored microbial tryptophan metabolism in irinotecan-treated rats, which was characterized by the enrichment of *Limosilactobacillus reuteri*, and elevated levels of indole-3-ethanol (IE) and indole-3-propionic acid (IPA). Multi-omics analysis further revealed a positive correlation between *L. reuteri* and IE and IPA. Consistent with this, DKT promoted *L. reuteri* proliferation, leading to the conversion of tryptophan to IE and IPA, which improved epithelial barrier damage in the irinotecan-treated Caco-2 cells. In addition, DKT suppressed the growth of Loop 1 β-glucuronidase (β-GUS)-producing bacteria, such as *Escherichia coli*. Furthermore, the main constituents of DKT selectively inhibited Loop 1 β-GUS activity independent of the gut microbiota, which reduced the intra-luminal level of 7-ethyl-10-hydroxycamptothecin, the toxic metabolite of irinotecan. Taken together, this study reveals a dual gut microbiota–driven mechanism by which DKT mitigates irinotecan–induced enterotoxicity, which provides a promising strategy for managing chemotherapy-related enterotoxicity.

## 1. Introduction

Irinotecan (CPT-11), a potent antineoplastic agent, is first-line treatment for advanced colorectal cancer (CRC), and is often combined with other chemotherapy drugs against CRC, small cell lung cancer, pancreatic cancer, and gastric cancer. Global market size of irinotecan hydrochloride injection reached 16.482 billion yuan (RMB) by 2024, and is estimated to reach 27.876 billion yuan (RMB) by 2030 (www.globalmarketmonitor.com.cn). Nevertheless, irinotecan-induced enterotoxicity, which is characterized by diarrhea, nausea and vomiting, seriously limit its clinical application and efficacy^1^. The incidence rate of delayed-onset diarrhea, the dose-limiting toxicity of CPT-11, is 60-80%, and that of III-IV grade diarrhea is 15-40%^2,3^. Severe diarrhea may trigger life-threatening complications, thus forcing chemotherapy discontinuation and worsening prognosis. In clinical practice, the management of chemotherapy-induced diarrhea primarily relies on loperamide, an opioid receptor agonist that reduces gut motility, or on broad-spectrum antibiotics aimed at suppressing gut microbiota^4^. However, these interventions may give rise to additional complications and are associated with a high rate of treatment failure, thereby underscoring the clinical need for safer and more effective therapeutic strategies^4^.

The gut microbiota is a key factor underlying the heterogeneity in chemotherapy outcomes^5^. CPT-11 disrupts the gut microbial ecology, while gut microbiota in turn modulate the toxicity of CPT-11 by metabolizing the drug and endogenous small molecule metabolites^5–9^. Gut microbial β-glucuronidase (β-GUS) is produced by diverse bacterial phyla, and hydrolyzes the glucuronide conjugates of drugs that are formed in the liver into their active aglycone forms in the intestine^10^. CPT-11 undergoes sequential biotransformation in the liver to the inactive metabolite 7-ethyl-10-hydroxycamptothecin glucuronic acid (SN-38G), and is then excreted into the gut, where bacterial β-GUS reactivates SN-38G into the potent cytotoxin SN-38, thus contributing to enterotoxic effects^11^. Notably, the catalytic activity and substrate selectivity of the different β-GUS subtypes vary depending on their loop structure^12,13^. Loop 1 β-GUS exhibits high catalytic efficiency for SN-38G, and correlates with intestinal barrier injury and diarrhea severity^14–16^. Therefore, blockade of gut microbial β-GUS to limit intra-luminal SN-38 production represents a new therapeutic strategy for preventing irinotecan-induced enterotoxicity. However, monocolonization by β-GUS-producing bacteria cannot fully replicate irinotecan-associated intestinal damage, suggesting the involvement of β-GUS-independent mechanisms^17^. Gut microbial metabolites also influence intestinal homeostasis and chemotherapy-related injury. CPT-11 is known to disrupt multiple classes of microbial metabolites that are essential for intestinal homeostasis^18–23^. The tryptophan metabolism pathway is notably dysregulated in the intestine of patients who experience diarrhea following irinotecan treatment compared to the non-diarrhea patients. In particular, indole-3-acetate (IAA) impairs intestinal stem cell self-renewal and epithelial repair capacity through inhibiting PI3K-Akt signaling pathway, thus exacerbating intestinal toxicity^9^. Moreover, in a rat model of irinotecan-induced diarrhea, the severity of intestinal damage was significantly correlated with tryptophan-kynurenine (KYN) metabolic pathway^18^. Therefore, the precise role of microbiota-derived metabolism in irinotecan-induced enterotoxicity requires urgent clarification.

Traditional Chinese medicine formulations have been utilized in clinical practice for centuries with well-established safety and efficacy. Daikenchuto (DKT) is widely used to treat digestive system disorders. Our previous study identified 74 chemical compounds in DKT, among which the core representative constituents, including skimmianine (SMN), 6-shogaol (6-SGL) and ginsenoside Rb_1_ (GRb_1_), among others were recognized^24^. DKT exhibits multifaceted pharmacological activities including gut microbiota modulation^25,26^, anti-inflammatory effects^27^, and antineoplastic activities^28,29^. In our study, DKT mitigated both acute and chronic colitis by regulating gut microbiota-derived tryptophan metabolites and activating aryl hydrocarbon receptor (AhR) signaling pathway^24^. Moreover, DKT can inhibit irinotecan-induced intestinal mucosal inflammation and apoptosis, and preserve small intestinal integrity^30^. Altogether, these studies point that DKT can exert its protective effects against irinotecan-induced enterotoxicity via the regulation of gut bacterial metabolism. However, the precise mechanism underlying this metabolic remodeling remains largely unknown.

The aim of this study was to evaluate the protective potential of DKT on irinotecan-induced enterotoxicity and elucidate the underlying gut microbiota-driven mechanisms. This study integrates 16S rRNA sequencing analysis, tryptophan-targeted metabolomics, fecal microbiota transplantation, *in vitro* bacteria-drug co-culture systems, β-GUS activity assays, and *in vivo* evaluation of anti-tumor efficacy to elucidate a dual mechanism by which DKT enhances intestinal barrier function by targeting *Limosilactobacillus reuteri*-tryptophan axis, and reduced intra-luminal SN-38 accumulation through inhibition of *Escherichia coli* and its β-GUS activity, without compromising the antineoplastic efficacy of irinotecan.

## 2. Materials and methods

### 2.1. Reagents

DKT was obtained from YUBARI TSUMURA Co., Ltd. (Tokyo, Japan). Irinotecan Hydrochloride Injection was obtained from Qilu Pharmaceutical Co., Ltd. (Shandong, China). Loperamide hydrochloride capsules were obtained from Xi’an-Janssen Pharmaceutical Co., Ltd. (Xi’an, China). The following compounds were sourced from Aladdin Biochemical Technology Co., Ltd. (Shanghai, China): 4 kDa fluorescein isothiocyanate (FITC)-dextran, 4-nitrophenyl-β-D-glucopyranoside (PNPG), phosphate buffered saline buffer (PBS, pH 6.8), D-saccharic acid 1,4-lactone hydrate (DSA), L-Tryptophan (Trp), indole-3-ethanol (IE), and indole-3-propionic acid (IPA). The following compounds were sourced from Baoji Herbest Bio-Tech Co., Ltd. (Baoji, China): skimmianine (SMN), hydroxy-α-sanshool (HαS), hydroxy-β-sanshool (HβS), 6-shogaol (6-SGL), 6-gingerol (6-GGL), ginsenoside Rb_1_ (GRb_1_), ginsenoside Re (GRe), and ginsenoside Rg_1_ (GRg_1_).

### 2.2. Cell culture

Caco-2 cell line and CT26 cell line, purchased from the American Type Culture Collection (ATCC, Rockville, MD, USA), were cultured in DMEM or 1640 culture medium supplemented with 10% fetal bovine serum and 1% penicillin-streptomycin solution in an incubator with a humidified atmosphere of 5% CO_2_ at 37℃. All culture reagents are sourced from Sangon Biotech (Shanghai, China).The Caco-2 cell was seeded into six-well plates until reaching 80-90% confluence, and divided into the control, CPT-11(150 μM CPT-11), IPA (0.1 mM IPA plus 150 μM CPT-11), and IE group (0.1 mM IE plus 150 μM CPT-11), and subjected to the corresponding treatments for a 24-h duration. Then the Caco-2 cells were subsequently collected for the extraction of either RNA or protein.

### 2.3. Animal experiments

Specific pathogen free (SPF) Male Sprague Dawley rats (6-8 weeks, 180-200 g), male Balb/c-mice and Balb/c-nude mice (6-8 weeks, 18-22g) were obtained from the GemPharmatech Co., Ltd (Jiangsu, China) (SCXK (Su) 2023-0009). Animals were acclimatized to standard SPF environment for a week and allowed to feed and drink ad libitum. All animal procedures were approved by the Animal Ethical Council of Xi’an Jiaotong University Health Science Center (permit NO. XJTUAE2025-2677).

For establishing an irinotecan-induced enterotoxicity rat model and drug treatment, the rats were randomly divided into Control, Model, Loperamide (LOP, the positive control drug, 2 mg/kg), DKT-L (0.25 g/kg) and DKT-H (0.75 g/kg) (*n*=6 per group). Rats of LOP, DKT-L and DKT-H group were given daily gavage of corresponding dosage of LOP and DKT for a 3-day preintervention, and other groups were given daily gavage of drinking water. On Days 4-7, rats received irinotecan (75 mg/kg) via intraperitoneal injection for modeling except control group, and diarrhea scores were monitored during the whole experimental period according to the reported scoring criteria^31^: 0, normal; 1, slightly wet; 2, moderate wet and unformed stool; 3, watery stool. Rats were treated for consecutive 7 days and sacrificed on day 8. 1-cm segments from small intestine and distal colon were retained in 4% paraformaldehyde or Carnoy’s fixative for histological examination, periodic acid-schiff (PAS) staining, immunofluorescence analysis, and immunohistochemistry analysis respectively. At last, the remaining intestinal tissues and contents were frozen in liquid nitrogen and kept at -80°C freezer for later experiments.

Male Balb/c mice were randomly divided into DKT (1.1 g/kg), ABX-Model, ABX-DKT (1.1 g/kg) and ABX-DKT-FMT (0.1 mL/10 g) (*n*=6 per group). The pre intervention period and modeling period were the same as above. For pseudo-germ-free mice modeling, antibiotic cocktail (ABX, vancomycin hydrochloride 0.1 g/kg, ampicillin 0.2 g/kg, neomycin sulfate 0.2 g/kg, and metronidazole 0.2 g/kg, 0.1 mL/10 g) was orally administered to mice 7 days before DKT and DKT-FMT pre intervention. The feces of DKT group donor mice were diluted with physiological saline 1:10 (w/v) (g/mL). Then the suspension was centrifuged at 4℃ for 500 g for 3 minutes to obtain the supernatant. ABX-DKT-FMT group received fecal microbiota solution (0.1 mL/10 g) prepared from the donor DKT group.

For the tumor-bearing mice modeling, CT26 cells (1×10^7^ cells/mL) were subcutaneously injected into the right flank of Balb/c–nude mice. Mice were randomly divided into Control, CPT-11 and CPT-11+DKT group when the tumor volume reaches 50-100 mm^3^. CPT-11+DKT group was orally administered with high dosage of DKT on days 1-7, and CPT-11 and CPT-11+DKT groups were administered CPT–11 (60 mg/kg) via intraperitoneal injection on days 2-5. Tumor volume was calculated by measuring the largest diameter (*a*) and perpendicular (*b*) of the tumor (tumor volume = *a*×*b*^2^/2). Body weight and diarrhea scores after the last injection of irinotecan 48 hours were monitored according to the above criteria. Mice were sacrificed, and spleens and tumors were weighed on Day 8.

### 2.4. Enzyme-linked immunosorbent assay (ELISA)

Using commercial ELISA kits (Jianglai Biotechnology Co., Ltd., Shanghai, China), the levels of tumor necrosis factor (TNF–α), interleukin–1β (IL–1β), and interleukin–10 (IL–10) in both small intestine and colon tissues were measured, following the protocol provided by the manufacturer.

### 2.5. Histopathological evaluation

For hematoxylin and eosin (H&E) staining, small intestine, distal colon and tumor tissues were fixed in 4% paraformaldehyde fixative overnight, and then embedded and sectioned at 5 μm in accordance with standard procedures. For PAS staining, the small intestine and distal colon tissues were fixed in Carnoy’s fixative. Goblet cells were counted, and the length of villi and crypts were measured using Image J software.

### 2.6. Immunofluorescence staining

Paraffin slices of small intestine tissue were deparaffinized, rehydrated and blocked endogenous peroxidase activity. After blocking with 3% bovine serum albumin, they were incubated overnight at 4°C with primary antibodies. Following this, a Cy3-conjugated secondary antibody was applied for 50 minutes at room temperature. Finally, nuclei were stained using DAPI (Servicebio, Wuhan, China) before visualization on a laser scanning confocal microscope (Leica, Wetzlar, Germany).

### 2.7. Intestinal permeability measurement

The assessment of intestinal permeability was conducted as our previously established method^32^. Briefly, after a 4-hour fast, rats were orally gavage of 4 kDa FITC-dextran (600 mg/kg). Serum was separated 5h later and serum FITC-dextran levels were quantified using the standard curve with fluorescence detection at 485 nm excitation and 525 nm emission wavelength (BioTek, VT, USA).

### 2.8. Western blot analysis

The protein lysates of Caco-2 cell line were prepared using RIPA Lysis Buffer (Servicebio, Wuhan, China) plus protease and phosphatase inhibitors cocktail. After that, the total protein concentrations in RIPA Lysis buffer were determined with a BCA assay kit (Servicebio, Wuhan, China). Western blot analysis was conducted as detailed in previous study^32^. The primary antibodies included Cannabinoid receptor 1 (CB1, 1:2000), ZO-1 (1:5000), Occludin (1:8000), and GAPDH (1:10000). All antibodies are sourced from Proteintech (Wuhan, China). Protein bands were subsequently visualized using a GS-700 imaging densitometer (Bio-Rad, Hercules, CA, USA).

### 2.9. Reverse transcription quantitative polymerase chain reaction (RT-qPCR)

TRIzol Reagent was used to extracted total RNA from the small intestine tissues and Caco-2 cell line. RNA was reverse transcribed to cDNA using the Evo M-MLV RT premix, and quantitative PCR was performed with the SYBR Green Premix Pro Taq HS qPCR Kit. All reagents used for RT-qPCR are sourced from Accurate Biotechnology Co., Ltd. (Hunan, China). The amplification program consisted of an initial denaturation cycle at 95°C for 2 min, followed by 40 cycles of denaturation at 95°C for 5 s, annealing at 60°C for 30 s. Transcriptional differences (*Tjp1*, *Ocln*, *Cldn1*, *Cldn4*, *Cnr1* and *Cnr2* in small intestine tissues of rats and *TNF*, *IL6*, *IL1B*, *TJP1*, *OCLN*, *CLDN1* and *CNR1* in Caco-2 cell line) were analyzed using GAPDH as an internal control. Primer sequences are listed in supporting information (Table S1).

### 2.10. 16S rRNA sequencing analysis

Microbial genomic DNA was extracted from small intestine contents using the E.Z.N.A.® soil DNA Kit (Omega Bio-tek, Norcross, GA, U.S.). The bacterial 16S rRNA genes were amplified using the universal bacterial primers 27F (5’-AGRGTTYGATYMTGGCTCAG-3’) and 1492R (5’-RGYTACCTTGTTACGACTT-3’). The PCR amplification was performed using T100 Thermal Cycler PCR thermocycler (Bio-Rad, CA, USA). The amplification program consisted of an initial denaturation at 95℃ for 3 min, followed by 27 cycles of denaturation at 95℃ for 30 s, annealing at 60°C for 30 s and extension at 72℃ for 45 s, and single extension at 72℃ for 10 min, and end at 4℃. Then the PCR products were purified and quantified. DNA library was constructed with the SMRTbell prep kit 3.0 (Pacifc Biosciences, CA, USA) and were sequenced on the Pacbio Sequel IIe System (Pacifc Biosciences, CA, USA) by Majorbio Bio-Pharm Technology Co. Ltd. (Shanghai, China). According to the data of the OTU analysis, Shannon index, principal coordinate analysis (PCoA) based on Bray-curtis dissimilarity and significant species difference analysis were conducted to compare the differences among the groups. A correlation between gut microbiota and indicators of enterotoxicity was considered to be statistically robust if the *P*-value less than 0.05 based on the spearman’s correlation analysis.

### 2.11. Targeted metabolomics analysis

For sample preprocessing, appropriate samples were mixed with 400 μL of cooled methanol-water (4:1, v/v) containing 0.1% formic acid, internal standard [2H5]-Trp, and 1 mM butylated hydroxytoluene. Small steel beads were added to mixture, which was transferred to Automatic High-Speed Sample Grinder (Wonbio-E, Shanghai, China) and ground at 45 Hz for 2 min after being stored in a -20℃ freezer for 2 min. The mixture was sonicated in ice bath and centrifuged for 300 μL of supernatant, which was evaporated to dryness. 200 μL of cooled methanol-water with the same content was added to the residue. Repeat the above steps to obtain the dried residue, which was reconstituted with 200 μL of water (containing L-2-chlorophenylalanine) and sonicated. The mixture was centrifuged, and 200 μL of the supernatant was collected and filtered using a 0.22 μm hydrophilic filter membrane for LC-MS analysis. Additionally, quality control samples were prepared by mixing equal volumes of the extraction solutions from each sample.

Targeted metabolomics was carried out using UPLC-ESI-MS/MS (AB Sciex Qtrap 6500+/AB ExionLC). The metabolomics method was conducted using an ACQUITY UPLC HSS PFP (100 mm×2.1 mm, 1.8 μm) with a flow rate of 0.3 mL/min and the column temperature maintained at 40°C. Mobile phase A was water with 0.1% formic acid (v/v), and B was methanol with 0.1% formic acid (v/v). The gradient-elution procedure was as follows: 0 min A/B (99:1, v/v), 1 min A/B (99:1, v/v), 6min A/B (5:95, v/v), 7 min A/B (5:95, v/v), 7.01 min A/B (99:1, v/v), 8min A/B (99:1, v/v). The injection volume was 5 μL. Table S2 in the supporting information provides the mass spectrometry parameters.

### 2.12. Bacterial β-GUS activity detection

β-GUS activity in intestinal contents was assessed by PNPG assay. 100 mg frozen intestinal contents were homogenized in 300 μL PBS (pH 6.8) and then centrifuged to collect the supernatant of all samples. The total protein concentration was then measured with the BCA Protein Assay Kit (Servicebio, Wuhan, China). Fecal supernatants containing 100 μg/ml protein were incubated with 0.8 mM PNPG and PBS for 40 min at 37°C. Finally, the reaction was terminated by sodium hydroxide (0.5 M) and the absorbance was read at 405 nm.

Next, we investigated the effect of DKT components on Loop 1 β-GUS activity derived from *E. coli* (EcGUS) and β-GUS derived from bovine liver (BLGUS). DSA, 6-SGL, 6-GGL, HαS, HβS, SMN, GRb_1_, GRe, and GRg_1_ were prepared as working solution (1 mM) using dimethyl sulfoxide solution (DMSO). EcGUS (37.5 U/mL) and BLGUS (1 KU/mL) were prepared as working solution using PBS. For the activity assay of EcGUS, the final incubation system included 10 uL EcGUS (final concentration: 3.75 U/mL), 10 uL components/DSA/DMSO (100 μM), and 10 uL PNPG (250 μM) and 70 uL PBS. For the activity assay of BLGUS, the final incubation system included 20 uL BLGUS (200 U/mL), 10 uL components/DSA/DMSO (100 μM), and 20 uL PNPG (500 μM) and 50 uL PBS. After a 40-minute incubation of EcGUS and a 3-h incubation of BLGUS at 37°C, reaction was terminated by sodium hydroxide (0.5 M) and the absorbance was read at 405 nm. DMSO was used as a blank control in this experiment and all experiments were repeated for three times.

To detect IC_50_ value on the EcGUS activity of DSA, 6-SGL, HαS and HβS, positive inhibitor DSA and three components were prepared into different concentrations working solutions ranging from 0-1000 uM. The mixture contained EcGUS (3.75 U/mL), PNPG (250 μM), and inhibitor solution at various concentrations. Each concentration was tested in triplicate to calculate the relative activity. Finally, IC_50_ values were calculated by plotting the concentration response curve.

### 2.13. Determination of SN-38 content in intestine contents

1 g feces per group were mixed with 100 uL DMSO and 900 uL of methanol and processed by vortexing and sonication for 5 min, followed by centrifugation to obtain the supernatants. The supernatant was filtered using a 0.22 μm filter membrane. Analysis was subsequently performed on an Agilent® 1100 LC-MS system using an Agilent ZORBAX SB -C18 column (5 μm, 4.6 mm × 250 mm) with a gradient elution program. The mobile phase consisted of an aqueous solution containing 0.1% formic acid (A) and pure acetonitrile (B). Under column temperature conditions of 40°C, the injection volume was 10 μL, and the flow rate was set at 1.0 mL/min. The gradient elution was as follows: 10% B → 10% B (0-3 min), 10% B → 35% B (3-6 min), 35% B → 65% B (6-18 min), 65% B → 95% B (18-21 min) and 95% B → 10% B (25-28 min).

### 2.14. Intestinal bacterial growth assay

To investigate the effects of DKT, SMN, HαS, HβS, 6-SGL, 6-GGL, GRb_1_, GRe, and GRg_1_ on the growth of *L. reuteri* ATCC 23272 and two strain of *E. coli* ATCC 25922 and 35218, DKT (final concentrations: 1, 2, and 8 mg/mL) and 6-SGL, 6-GGL, HαS, HβS, SMN, GRb_1_, GRe, and GRg_1_ (final concentrations: 200 μM) were added to Man, Rogosa, and Sharpe (MRS) Broth culture medium. OD_600_ values were measured from 0 to 24 hours respectively. The growth curves of *L. reuteri* and two strain of *E. coli* were calculated using GraphPad Prism 8.0.

### 2.15. IPA and IE concentration detection in culture supernatant of Limosilactobacillus reuteri

Appropriate volumes of high and low concentration tryptophan and normal MRS culture medium were prepared, including Trp- (MRS medium), Trp+ (MRS medium plus 0.1 mM tryptophan), and Trp++ (MRS medium plus 1 mM tryptophan), respectively. 1 mL bacterial solution of *L. reuteri* was added to 14 mL MRS medium containing different concentrations of Trp and incubated anaerobically at 37℃ for 24 hours. 1 mL suspension was centrifuged at each time point of 0, 6, 12, and 24 hours, to collect the supernatant. IE and IPA in supernatant of culture medium were determined utilizing specific ELISA kits (Youxuan Biotechnology Co., Ltd., Shanghai, China) in accordance with the instructions provide by manufacturer.

### 2.16. Statistics analysis

Data are presented as the mean ± standard error of the mean (SEM) and were analyzed using GraphPad Prism 8.0 (GraphPad Software, La Jolla, CA). Statistical differences were conducted using two-tailed unpaired Student’s *t*-test, or one-way or two-way analysis of variance (ANOVA) followed by Dunnett’s *post hoc* multiple comparison’s test. The gut microbiota sequencing data were analyzed on the online platform of Majorbio Cloud Platform (www.majorbio.com), performing a two-tailed Wilcoxon rank-sum test. The corresponding statistical differences are indicated as follows: \**P* < 0.05, \*\**P* < 0.01, \*\*\**P* < 0.001 *vs*. Control group, ^#^*P* < 0.05, ^##^*P* < 0.01, ^###^*P* < 0.001 *vs*. Model group.

## 3. Results

### 3.1 DKT alleviated pathological symptoms of irinotecan-induced enterotoxicity

An irinotecan-induced enterotoxicity model was established by administering irinotecan via intraperitoneal injection for 4-7 days (Fig. 1A). By 24 h after the first injection, the rats exhibited a sharp decrease in body weight and experienced delayed-onset diarrhea. DKT was orally administered to the rats once daily for seven consecutive days at two different dosages (0.25 and 0.75 g/kg/day), and the higher dose effectively mitigated weight loss and diarrhea (Fig. 1B-C). Although loperamide, a first–line agent for chemotherapy–induced diarrhea, also alleviated the symptoms of diarrhea, it failed to prevent body weight loss during the experimental period. Pathological analysis of small intestine specimens revealed that DKT administration attenuated crypt destruction, villous shortening, and inflammatory cell infiltration induced by irinotecan (Fig. 1D-F). In addition, DKT also reversed the elevation in the pro-inflammatory cytokines TNF-α and IL-1β (Fig. 1G), indicating that it can mitigate irinotecan-induced inflammation. However, no significant differences in colon length or the colon weight–to–length ratio were observed among groups. Furthermore, neither significant damage nor inflammatory responses were detected in the colon of irinotecan-induced enterotoxicity rats (Fig. S1A-C). Taken together, DKT alleviated disease symptoms, histopathological damage, and inflammatory responses associated with irinotecan-induced enterotoxicity in a rat model.

**Figure 1.**
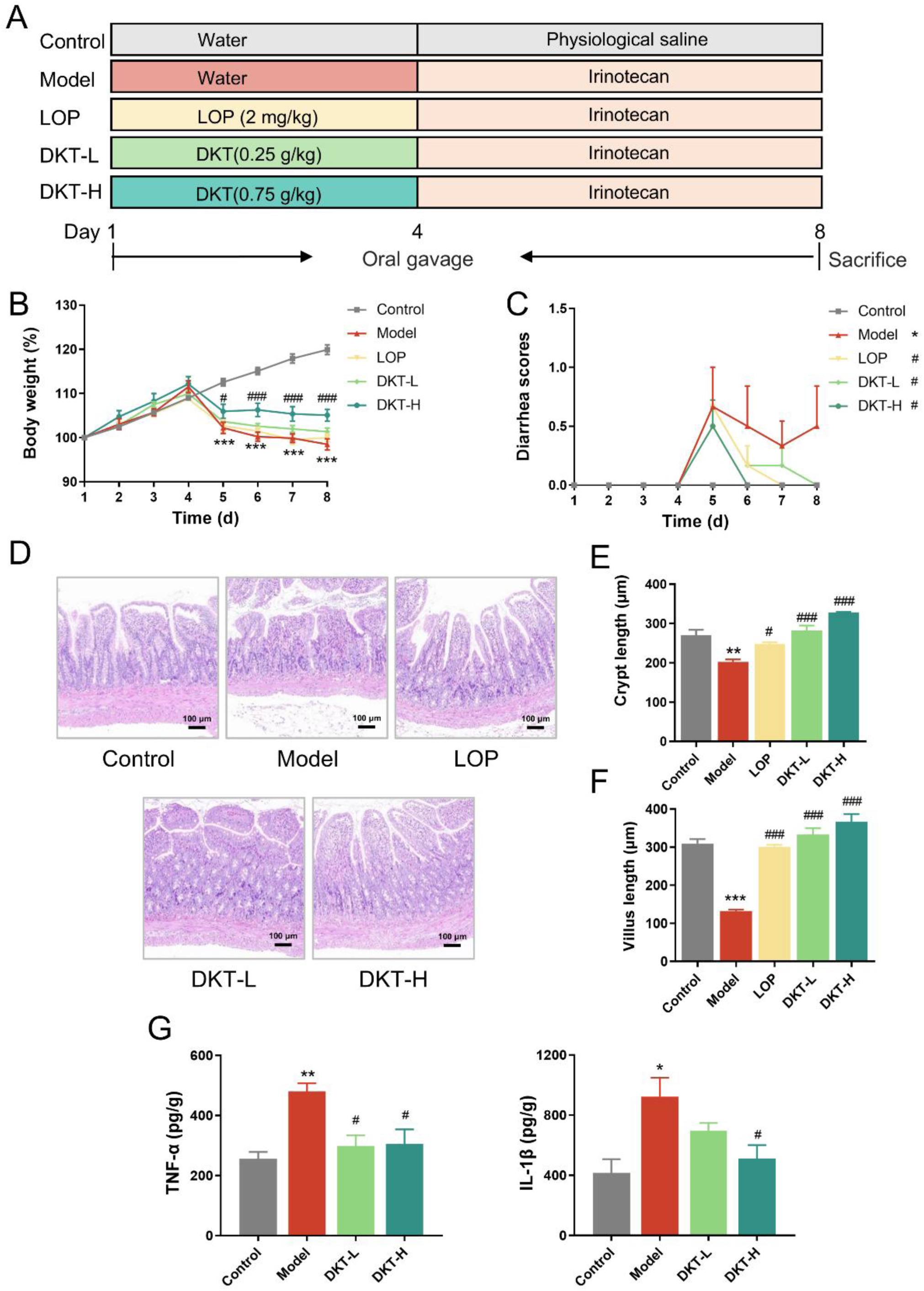
DKT alleviated pathological symptoms of irinotecan-induced enterotoxicity. (A) Experimental scheme. DKT at dosage of 0.25 g/kg/d (DKT-L) and 0.75 g/kg/d (DKT-H), Loperamide (LOP) at dosage of (2 mg/kg/d) were orally administered daily to rats during the enterotoxicity process. Enterotoxicity rats was induced by intraperitoneal injection of irinotecan (75 mg/kg) during days 4-7. (B) Percentage of body weight during the experiment (*n*=6). (C) Diarrhea scores during the experiment (*n*=6). (D) Representative hematoxylin and eosin (H&E) stained pictures of the small intestine (scale bars = 100 μm). (E) Crypt length in small intestine tissues (*n*=3). (F) Villus length in small intestine tissues (*n*=3). (G) Inflammatory cytokine levels (TNF-α and IL-1β) in small intestine tissues (*n*=3). Data are presented as the mean ± SEM. \**P* < 0.05, \*\**P* < 0.01 and \*\*\**P* < 0.001 *vs*. Control group, ^#^*P* < 0.05 and ^###^*P* < 0.001 *vs*. Model group.

### 3.2 DKT restored intestinal barrier function and integrity in the irinotecan-treated rats

Intestinal barrier disruption is a central pathological event in irinotecan–induced enterotoxicity, serving as both a critical driver of mucosal inflammation and a key determinant of disease severity^33,34^. Therefore, we assessed intestinal barrier function among three groups. The serum level of FITC-dextran, a well–established indicator of intestinal permeability, was markedly elevated in the irinotecan–treated rats and effectively normalized by DKT (Fig. 2A), indicating that DKT restored the intestinal barrier function. This functional improvement was accompanied by cellular and molecular alterations. DKT treatment reversed the loss of goblet cells in the model group, which synthesize and secrete mucins to form a protective mucosal barrier^35^(Fig. 2B-C). Consistent with this, DKT upregulated the transcripts of tight junction protein-encoding genes, including *Tjp1*, *Ocln*, *Cldn1*, and *Cldn4* (Fig. 2D). Immunofluorescence staining further demonstrated that ZO–1 and Claudin–4 expression were notably diminished in the damaged small intestine of the model group. DKT administration normalized their expression to control levels (Fig. 2E-F). Overall, DKT effectively restored the intestinal barrier function and integrity in rats with irinotecan-induced enterotoxicity by preserving the goblet cells and enhancing tight junction protein expression.

**Figure 2.**
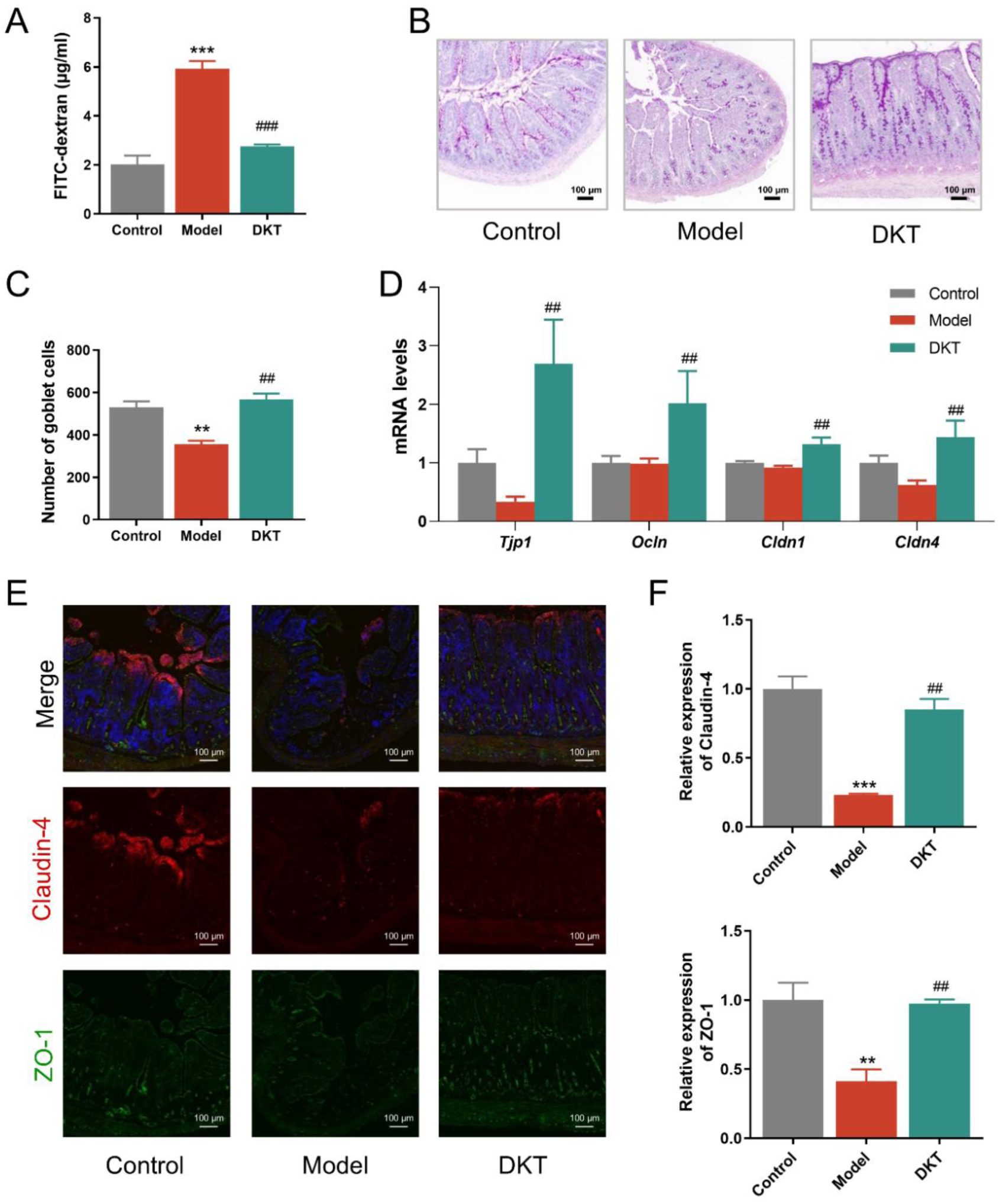
DKT restored intestinal barrier function and integrity in the irinotecan-treated rats. (A) The level of FITC-dextran in serum of different groups (*n*=3). (B, C) Representative periodic acid-schiff (PAS) staining of small intestine tissue sections (scale bars = 100 μm) (B) and the number of goblet cells (C). (D) The mRNA levels of tight junction (TJ) proteins *Tjp1*, *Ocln*, *Cldn1*, and *Cldn4*. (E) Representative immunofluorescence staining of the small intestine tissues for ZO-1 and Claudin-4. (F) Relative quantification of ZO-1 and Claudin-4 in the small intestine tissues in each group. Data are presented as the mean ± SEM (*n*=3). \*\**P* < 0.01 and \*\*\**P* < 0.001 *vs*. Control group, ^##^*P* < 0.01 and ^###^*P* < 0.001 *vs*. Model group.

### 3.3 DKT regulated the gut microbiota disorder associated with irinotecan-induced enterotoxicity

Gut microbiota critically contributes to the pathogenesis of irinotecan-induced enterotoxicity. Given the modulation of DKT on gut bacteria^25,26^, we analyzed the microbiota composition of the small intestine contents in the different groups through 16S rRNA sequencing. The model group exhibited markedly reduced microbial diversity, as reflected by the Shannon index (Fig. 3A). The control and model groups displayed distinct, non-overlapping clustering of microbial communities (Fig. 3B), indicating that irinotecan induced considerable gut microbial changes. In contrast, treatment with DKT significantly reversed irinotecan-induced gut dysbiosis by enhancing α-diversity and shifting the microbiota composition closer to normal mice. At the genus level, irinotecan markedly reduced *Limosilactobacillus* abundance but increased *Lactobacillus* abundance. (Fig. 3C). The relative abundance of *Limosilactobacillus* was subsequently restored by DKT, while that of *Lactobacillus* remained unaffected. Furthermore, heatmap analysis revealed significant correlation between *Limosilactobacillus* and key enterotoxicity phenotypes, including body weight, inflammatory factors, and tight junction proteins (Fig. 3D). We also profiled its microbiota composition at the species level, and observed a pronounced downregulation in the abundance of *L. reuteri*, *L. vaginalis*, *L. portuensis*, and *L. oris* in the model group (Fig. 3E). Notably, DKT treatment significantly restored *L. reuteri* and *L. vaginalis* abundance, which are involved in maintaining intestinal metabolism and homeostasis^36,37^. Collectively, these results demonstrate that DKT can effectively reverse the gut dysbiosis related to enterotoxicity by enriching beneficial bacteria like *L. reuteri* and *L. vaginalis*.

**Figure 3.**
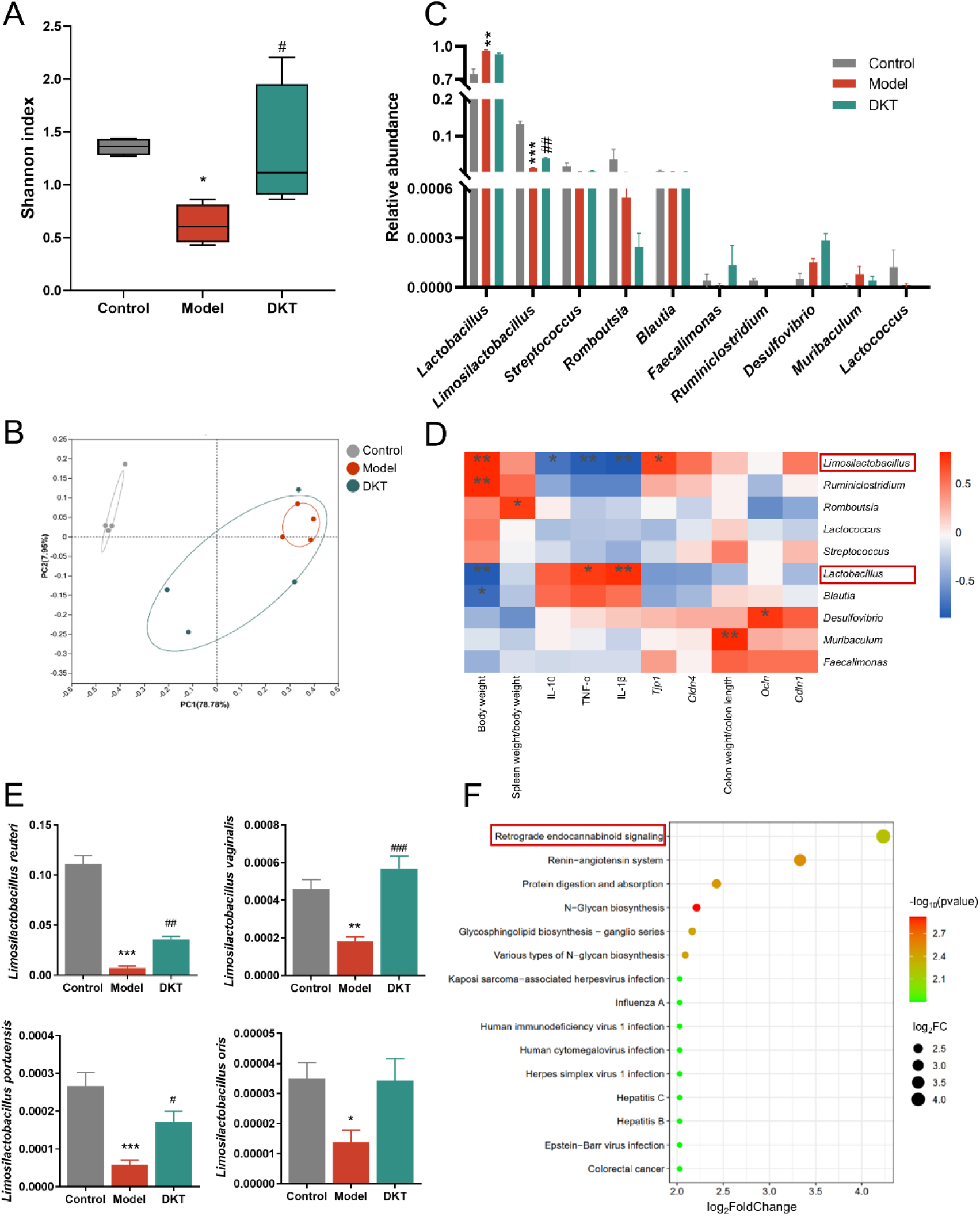
DKT regulated the gut microbiota disorder associated with irinotecan-induced enterotoxicity. (A) Alpha diversity analysis of gut bacterial diversity (Shannon index) from different rat groups (*n*=4). (B) Bray–Curtis principal coordinate analysis (PCoA) analysis of gut microbiota among three groups (*n*=4). (C) Relative abundance of dominant bacteria at the genus level in intestinal contents of different groups (*n*=4). (D) Heatmap analysis based on spearman correlation. \**P* < 0.05, \*\**P* < 0.01 and \*\*\**P* < 0.001. (E) Relative abundance of *Limosilactobacillus reuteri*, *Limosilactobacillus vaginalis*, *Limosilactobacillus portuensis*, *and Limosilactobacillus oris* in the intestinal contents from different groups (*n*=4). (F) Top 15 enriched pathways based on KEGG analysis between Model and DKT group. Data are presented as the mean ± SEM. \**P* < 0.05, \*\**P* < 0.01 and \*\*\**P* < 0.001 *vs*. Control group, ^#^*P* < 0.05, ^##^*P* < 0.01 and ^###^*P* < 0.001 *vs*. Model group.

To predict the functional implications of bacterial changes, we performed Kyoto Encyclopedia of Genes and Genomes (KEGG) enrichment analysis and visualized the top 15 pathways (Fig. 3F). Endocannabinoid signaling pathway was identified as the primary candidate mediating the therapeutic effect of DKT. The endocannabinoid system serves as a crucial interface between the host and the gut microbiota by regulating intestinal permeability, immune environment, secretory functions, and motility to maintain gut homeostasis^38–40^. Therefore, the key endocannabinoid receptors expression was analyzed among different groups. As shown in Fig. S2A-B, the mRNA and protein of cannabinoid receptor 1 (CB1) were significantly upregulated in the model group, and subsequently normalized by DKT treatment. In contrast, CB2 mRNA levels showed no significant differences among groups. Notably, CB1 is predominantly localized to the intestinal epithelium and enteric nervous system in the gastrointestinal tract, where its activation increases intestinal permeability and downregulates tight junction proteins^41^. Thus, DKT may enhance intestinal barrier integrity through microbiota–dependent modulation of CB1 signaling.

To assess whether an intact gut microbiota is indispensable for the therapeutic efficacy of DKT, we ablated the gut bacteria using an antibiotic cocktail (ABX). Despite confirmed bacterial clearance (Fig. S3B), DKT was able to attenuate the major signs of enterotoxicity, including weight loss, histopathological changes, inflammation, and intestinal barrier damage (Fig. S3C-F). Furthermore, fecal microbiota transplantation from DKT-treated mice into recipient animals also produced a therapeutic benefit (Fig. S3C-F). Taken together, these findings suggest that DKT mitigated irinotecan-induced enterotoxicity in a partially microbiota-dependent manner.

### 3.4 DKT restored tryptophan metabolism in irinotecan-treated rats

Clinical and preclinical studies have underscored the pivotal involvement of tryptophan metabolites in the pathogenesis of irinotecan-induced enterotoxicity^9,18^. Furthermore, the gut tryptophan metabolite IAA, which is known to exacerbate irinotecan-induced enterotoxicity, has been identified as a predictive biomarker for chemotherapy-related enterotoxicity^9^. Therefore, we next performed targeted metabolomics to profile tryptophan metabolites in the small intestine contents of different groups. Partial least squares discriminant analysis (PLS-DA) revealed distinct clustering in the composition of tryptophan metabolites among three groups. Following treatment with DKT, the metabolite profile shifted markedly and exhibited closer clustering to that of the control group (Fig. 4A-B). Notably, DKT exerted a holistic regulatory effect on tryptophan metabolism, which was evident from the significant changes in the indole, kynurenine, and serotonin pathways (Fig. 4C-E). Most tryptophan metabolites, including indole-3-lactic acid (ILA), IAA, indole-3-acetyl-alanine (IAA-Ala), indole-3-acetyl-aspartate (IAA-Asp), kynurenic acid (KYNA), xanthurenic acid (XA), and serotonin (5-HT), were significantly elevated, whereas those of indole-3-carboxaldehyde (ICA), indole-3-propionic acid (IPA), and indole-3-ethanol (IE), were markedly reduced in the model group, and restored upon DKT intervention. Notably, the levels of ILA and XA increased more than threefold and positively correlated with the severity of enterotoxicity, suggesting their potential as biomarkers (Fig. 4B). Furthermore, *Limosilactobacillus* positively correlated with IPA and IE, but negatively correlated with ILA and XA, indicating that this genus may alleviate irinotecan-induced enterotoxicity by enhancing the production of IPA and IE (Fig. 4F). Taken together, DKT ameliorated tryptophan metabolism disorders via enrichment of indole metabolites, specifically IPA and IE, and this process is likely mediated by *Limosilactobacillus*.

**Figure 4.**
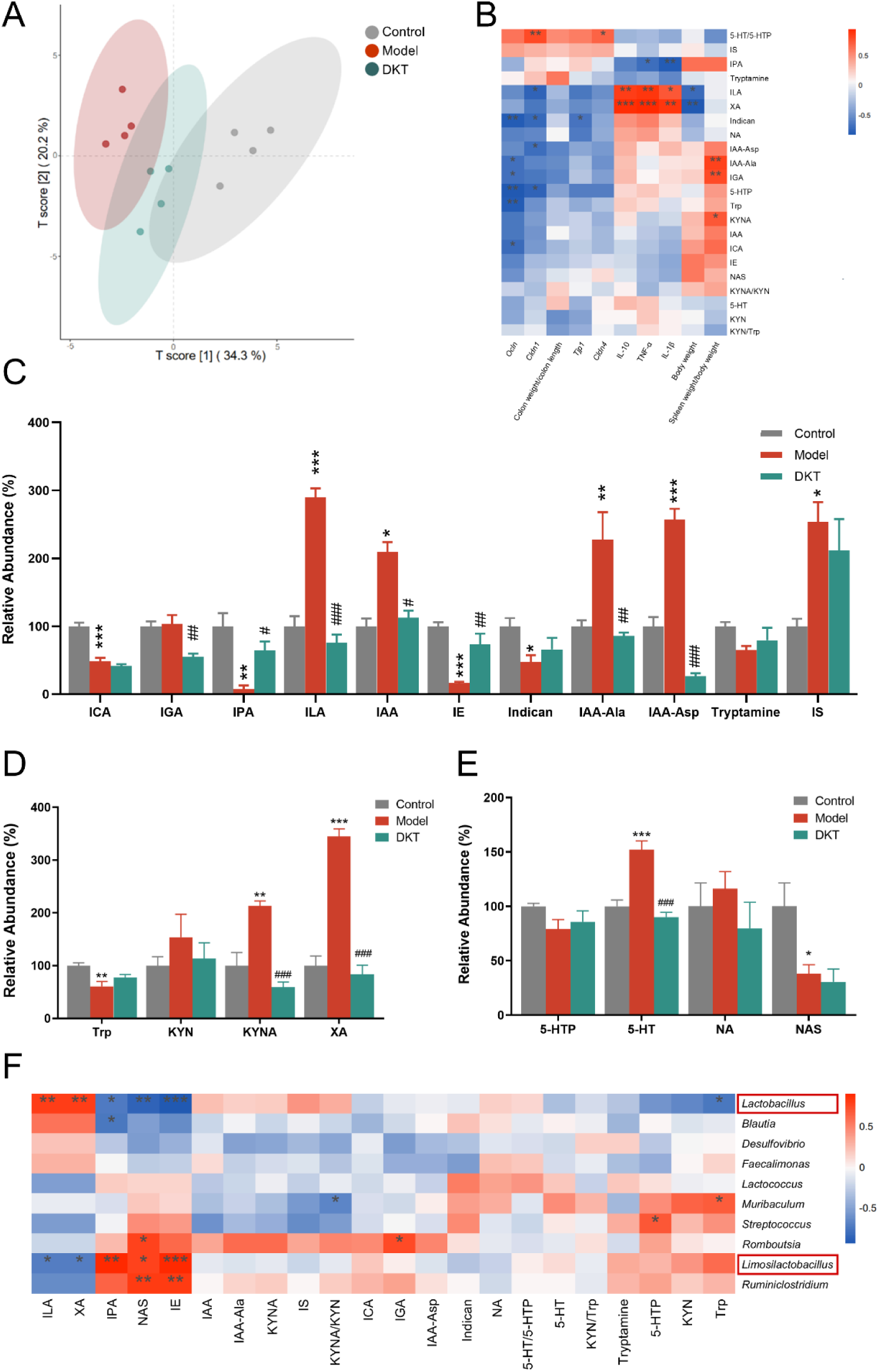
DKT restored tryptophan metabolism in irinotecan-treated rats. (A) Partial least-squares discriminant analysis (PLS-DA) among different groups. (B) Heatmap analysis based on spearman correlation between the enterotoxicity-related parameters and tryptophan metabolites in intestinal contents. \**P* < 0.05, \*\**P* < 0.01 and \*\*\**P* < 0.001. (C-E) The relative abundance of tryptophan metabolites in intestinal contents, including indole pathways (C), kynurenine pathway (D) and serotonin pathway (E). (F) Heatmap analysis based on spearman correlation between the tryptophan metabolites and dominant bacteria in intestinal contents. \**P* < 0.05, \*\**P* < 0.01 and \*\*\**P* < 0.001. Data are presented as the mean ± SEM (*n*=4). \**P* < 0.05, \*\**P* < 0.01 and \*\*\**P* < 0.001 *vs*. Control group, ^#^*P* < 0.05, ^##^*P* < 0.01 and ^###^*P* < 0.001 *vs*. Model group. ICA, indole-3-carboxaldehyde; IGA, indole-3-glyoxylicacid; IPA, indole-3-propionic acid; ILA, indole-3-lactic acid; IAA, indole-3-acetic acid; IE, indole-3-ethanol; IAA-Ala, indole-3-acetyl-alanine; IAA-Asp, indole-3-acetyl-aspartate; IS, indoxylsulfate; Trp, tryptophan; KYN, kynurenine; KYNA, kynurenic acid; XA, xanthurenic acid; 5-HTP, L-5-hydroxy tryptophan; 5-HT, serotonin; NA, nicotinicacid; NAS, N-acetyl-5-hydroxytryptamine.

### 3.5 Indole metabolites produced by L. reuteri mediated the entero-protective effect of DKT

Based on above results, we concluded that DKT mitigated enterotoxicity by enriching *Limosilactobacillus* indole metabolites, and repairing the intestinal barrier. Nevertheless, the complex interplay among these factors warrants further exploration. To evaluate the direct impact of DKT on bacterial growth, we incubated *L. reuteri* (the predominant enriched species) with DKT or each of its eight major components (SMN, HαS, HβS, 6-SGL, 6-GGL, GRb_1_, GRe, GRg_1_) *in vitro*. In contrast to its individual components, DKT promoted the growth of *L. reuteri*, implying that the effect arises from synergistic interactions among its multiple constituents, or from other pro-proliferative bioactive compounds, which was dependent on its concentration. These findings highlight the complex effects of herbal formulations on gut microbiota.

To assess the biosynthetic capacity of *L. reuteri* for IPA and IE, the bacterium was cultured in MRS supplemented with tryptophan (0.1 mM and 1 mM), and the levels of IPA and IE in the culture supernatant were quantified. Tryptophan supplementation significantly increased the content of these metabolites compared to that in the non-supplemented control. Furthermore, both metabolites accumulated in the supernatant in a time-dependent manner (Fig. 5B-C). This finding demonstrated that *L. reuteri* directly converted tryptophan into IPA and IE, which is consistent with previous reports^42,43^, and highlighted the potential role of these metabolites as the effectors of its probiotic function.

**Figure 5.**
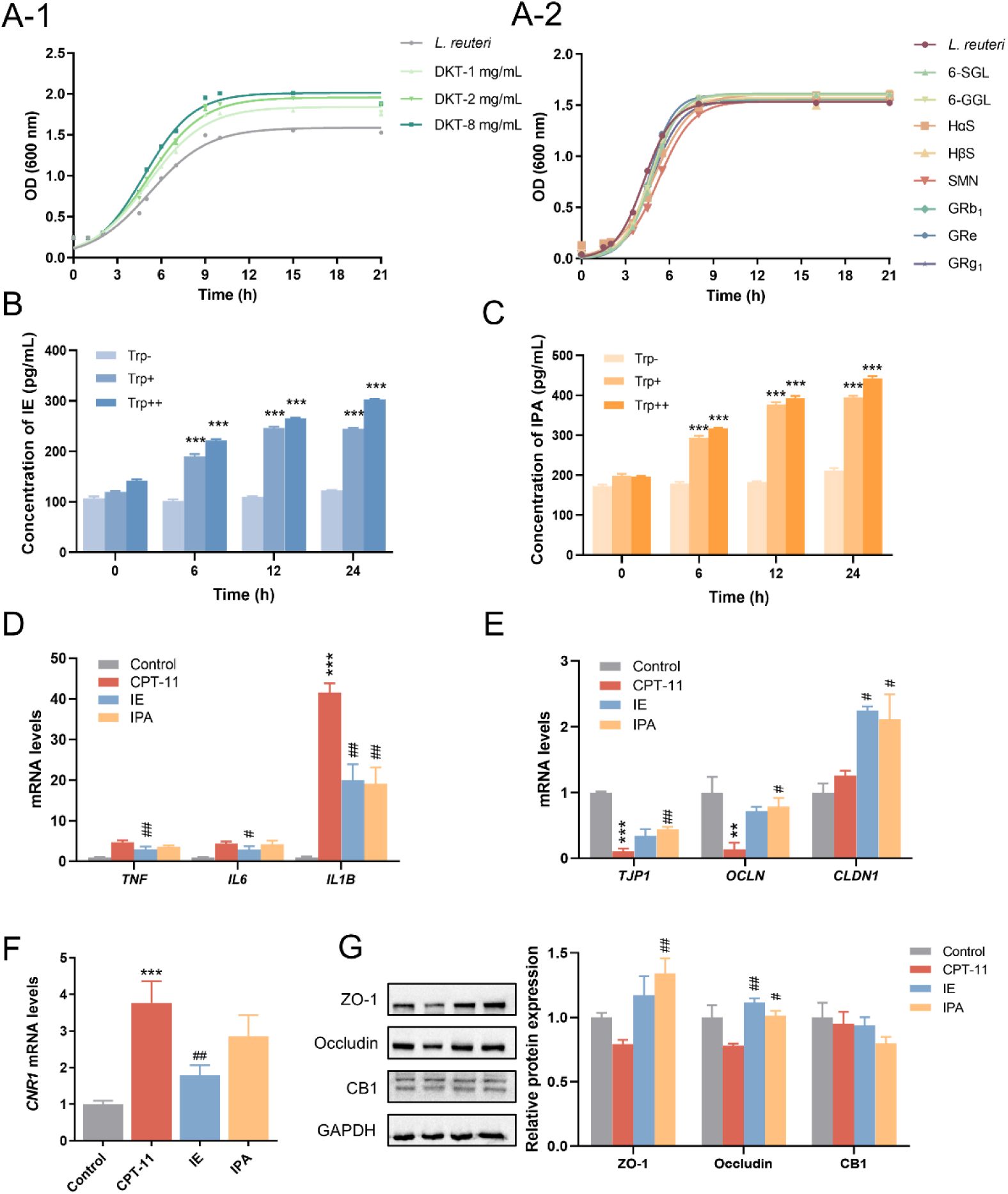
Indole metabolites produced by *L. reuteri* mediated the entero-protective effect of DKT. (A) Effect of DKT and its eight main components of DKT on the growth of *Limosilactobacillus reuteri*. (B, C) The effect of *Limosilactobacillus reuteri* on the conversion of Trp to IE (B) and IPA (C). \*\*\**P* < 0.001 vs. Trp-group. (D) The mRNA level of proinflammatory cytokines *TNF*, *IL6* and *IL1B* in Caco-2 cells. (E) The mRNA levels of tight junction (TJ) proteins *TJP1*, *OCLN*, and *CLDN1* in Caco-2 cells. (F) The mRNA levels of *CNR1* in Caco-2 cells. (G) Representative immunoblots and the relative expression levels of ZO-1, Occludin and CB1 proteins. Data are presented as the mean ± SEM (*n*=3). \*\**P* < 0.01 and \*\*\**P* < 0.001 *vs*. Control group, ^#^*P* < 0.05 and ^##^*P* < 0.01 *vs*. CPT-11 group.

A Caco-2 cell model of enterotoxicity was established by exposure to 150 µM irinotecan. Cells received 0.1 mM IE or IPA for a 24–hour incubation to evaluate their protective effects. IE significantly downregulated the transcripts of pro-inflammatory factors including *TNF*, *IL6*, and *IL1B* (Fig. 5D). Furthermore, irinotecan also downregulated the tight junction genes *TJP1* and *OCLN*. Both metabolites exhibited restorative effects, with IE upregulating *OCLN* and *CLDN1*, and IPA elevating *TJP1* and *CLDN1* mRNA levels (Fig. 5E). Consistent with this, both compounds upregulated ZO-1 and Occludin proteins expression (Fig. 5G), supporting their role in reinforcing intestinal barrier integrity. However, although irinotecan induction significantly increased *CNR1* mRNA, this change was not accompanied by a corresponding alteration in its protein expression (Fig. 5F). This suggests that DKT regulates CB1 and tight junction proteins in animal models through multiple pathways, and the indole metabolites IPA and IE are likely the major, though not exclusive, mediators in this process. Overall, our findings demonstrate that DKT mitigates irinotecan-induced enterotoxicity by promoting the proliferation of *L. reuteri*, which enhances intestinal barrier integrity by increasing IPA and IE levels.

### 3.6 DKT inhibited the growth of E. coli and its Loop 1 β-GUS activity to mitigate enterotoxicity

Microbial Loop 1 β–GUS, particularly that produced by *E. coli*, *Faecalibacterium prausnitzii*, and *Enterococcus faecalis*, catalyzes the SN–38G to the potent cytotoxin SN–38^10,34,44^. Notably, irinotecan intervention increased *E. coli* abundance by approximately 21-fold compared with the control group, making it the dominant species. DKT significantly reduced the abundance of three active β-GUS-producing bacteria, and restored *E. coli* levels comparable to those in the control group (Fig. 6A, Fig. S4A-B). Consistent with this, DKT significantly inhibited β-GUS activity in the small intestine contents (Fig. 6B), and lowered the intestinal concentration of SN-38 (Fig. 6C). This supports a mechanism wherein DKT limits SN-38 accumulation via dual inhibition of β-GUS activity and β-GUS-producing bacteria. However, the inhibitory effects of DKT were negligible in the colon (Fig. S1D).

**Figure 6.**
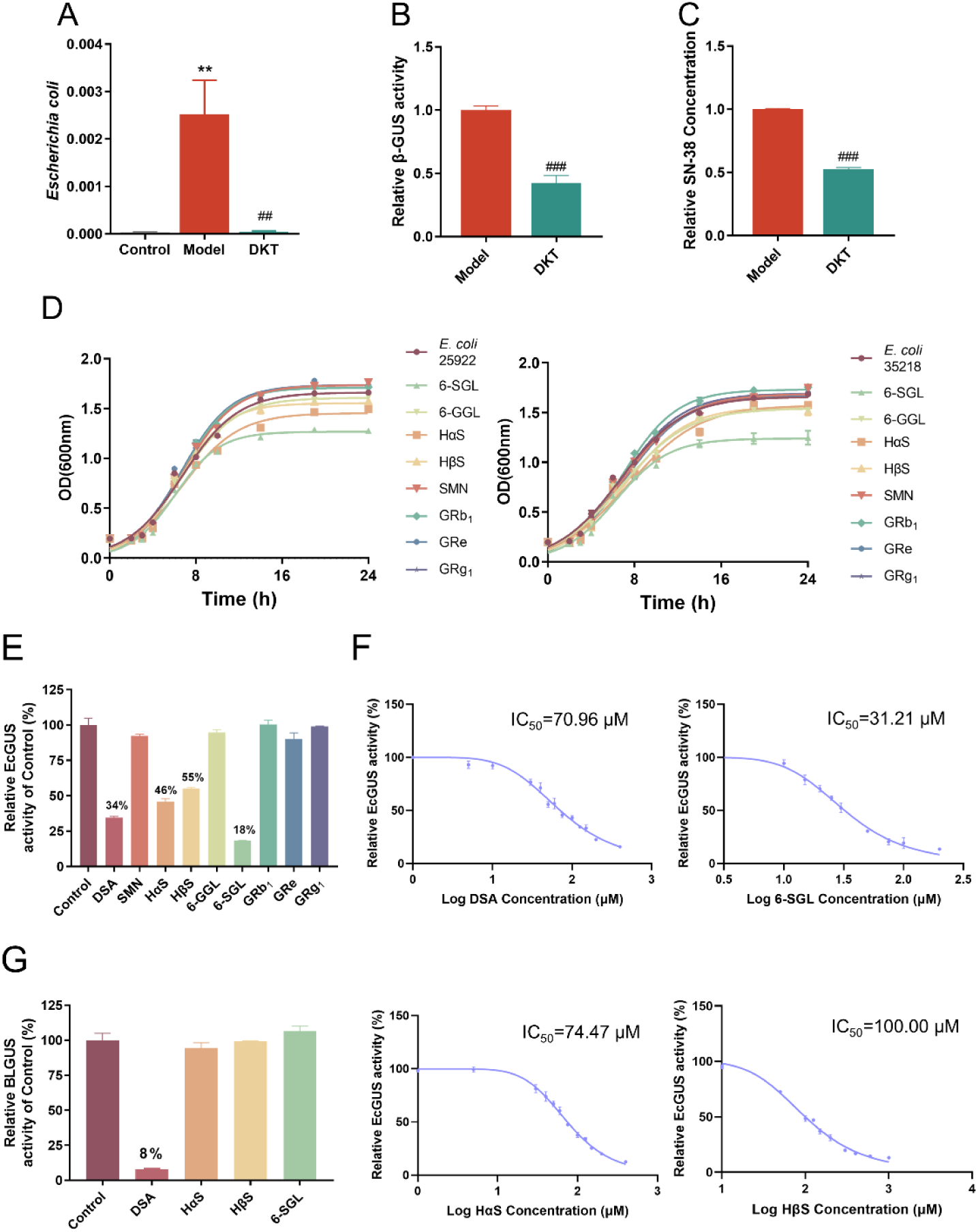
DKT inhibited the growth of *E. coli* and its Loop 1 β-GUS activity to mitigate enterotoxicity. (A) The relative abundance of *Escherichia coli* (*n*=4). (B) β-GUS activity in intestinal contents of Model and DKT groups (*n*=4). (C) The levels of SN-38 in intestinal contents of Model and DKT groups (*n*=3). (D) Effect of the eight main components of DKT on the growth of two strains of *E. coli* (*n*=3, ATCC: 25922 and 35218). (E, F) Effect of the eight main components of DKT on EcGUS activity (E) and its IC_50_ value (F) (*n*=3). (G) Effect of the eight main components of DKT on BLGUS activity (*n*=3). Data are presented as the mean ± SEM. \*\**P* < 0.01 *vs*. Control group, ^##^*P* < 0.01 and ^###^*P* < 0.001 *vs*. Model group.

Given that *E. coli* was the predominant Loop 1 β-GUS-producing species, we assessed the effects of eight key components on its growth profile over a period of 24 h. As shown in Fig. 6D, HαS, HβS, 6-SGL and 6-GGL inhibited the growth of the *E. coli* strains ATCC 25922 and ATCC 35218 by varying degrees. We further investigated the effects of these components (100 μM each) on the activity of Loop 1 β-GUS derived from *E. coli* (EcGUS), using DSA as the positive control^45^. DSA reduced EcGUS activity to 34% of the control, with an IC_50_ of 70.96 μM, whereas HαS, HβS, and 6-SGL inhibited the activity to 46%, 55%, and 18%, with respective IC_50_ values of 74.47 μM, 100.00 μM, and 31.21 μM (Fig. 6E-F). Notably, 6-SGL, a component derived from dried ginger, exhibited a stronger inhibitory effect than DSA. Given the broad distribution of β-GUS in mammalian tissues, non-selective inhibitors may disrupt the normal physiological processes in mammals^46^. Consequently, we evaluated the selectivity of DSA, HαS, HβS and 6-SGL against β-GUS activity in bovine liver (BLGUS). As shown in Fig. 6G, the inhibitory effect of DSA on mammalian β-GUS was 8%, which still carries the risk of disrupting host metabolic processes. In contrast, HαS, HβS and 6-SGL exhibited negligible effects on mammalian β-GUS activity. We also analyzed the effects of four tryptophan metabolites, including IE, IPA, ILA, and XA, on β-GUS activity. As shown in Fig. S4C and D, all tested metabolites had negligible inhibitory effects on the activities of EcGUS and BLGUS. This suggests that DKT reduces SN-38 concentration primarily through the direct regulation of β-GUS-producing bacteria and inhibition of β-GUS activity, rather than via an indirect effect mediated by microbial metabolism.

In summary, DKT alleviates irinotecan-induced enterotoxicity by suppressing active β-GUS-producing bacteria and specifically inhibiting Loop 1 β-GUS derived from *E. coli*. The DKT component 6-SGL exerted the most potent inhibitory effect, indicating that it is the key pharmacological agent underlying the protective effects of DKT.

### 3.7 DKT mitigated irinotecan–induced enterotoxicity without compromising anti-tumor efficacy

Since DKT reduced intra-luminal SN-38 levels in rats with irinotecan-induced enterotoxicity, and SN-38 is the key metabolite with anti-tumor activity, we hypothesized that DKT might interfere with the therapeutic effect of CPT-11. Consequently, we assessed whether DKT affects the anti-tumor activity of irinotecan. Balb/c-nude mice were subcutaneously inoculated with CT26 colon cancer cells for tumor-bearing mice modeling, and subsequently treated with CPT-11 and DKT (Fig. 7A). CPT-11 monotherapy led to significant weight loss and splenic atrophy, and the mice developed severe diarrhea 48 h after the final injection. DKT intervention markedly attenuated these pathological effects, corresponding to a reduction in CPT-11-associated toxicity (Fig. 7B-C). Furthermore, CPT-11 monotherapy potently inhibited tumor growth, and the combination therapy with DKT did not interfere with its anti-tumor activity (Fig. 7D-F). Furthermore, CPT-11 monotherapy and its combination with DKT led to the disintegration of tumor tissue structure and increased apoptosis rates in tumor cells (Fig. 7G-H). Overall, these results suggest that DKT effectively reduces the enterotoxicity of CPT-11 without compromising its anti-tumor activity.

**Figure 7.**
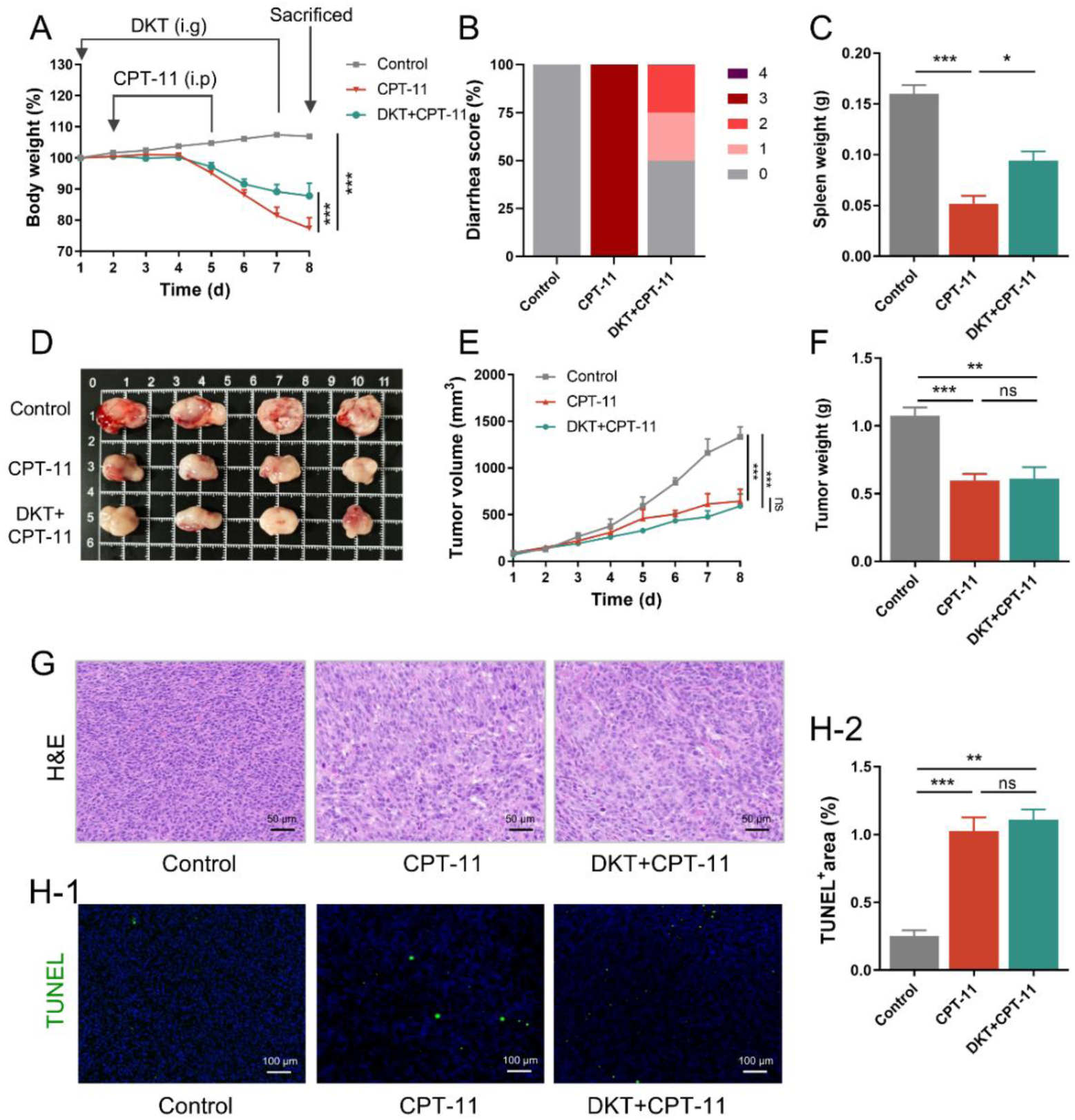
DKT mitigated irinotecan–induced enterotoxicity without compromising anti-tumor efficacy. (A) Tumor-bearing mice were treated with 60 mg/kg CPT-11 (i.p) on Days 2-5 and 1.1 g/kg DKT (i.g) on Days 1-7. Body weight of tumor-bearing mice in each group. i.p, intraperitoneal injection. i.g, oral gavage. (B) Diarrhea scores of mice in each group 48 hours after the last injection of CPT-11 (*n*=4). (C) Spleen weight of mice in each group (*n*=4). (D) Gross images of the tumors in each group. (E) Tumor volume of mice in each group throughout the experimental process (*n*=4). (F) Tumor weight of mice in each group on Day 8 (*n*=4). (G) Representative H&E staining images of tumors in mice in each group (scale bars = 50 μm). (H) Representative TUNEL staining images and quantification of TUNEL-positive tumor cells of mice in each group (scale bars = 100 μm). Data are presented as the mean ± SEM. \**P* < 0.05, \*\**P* < 0.01 and \*\*\**P* < 0.001.

## 4. Discussion

The non-specific cytotoxicity of CPT-11 poses a major clinical challenge due to the significant intestinal epithelium and gut dysbiosis. Studies increasingly support the translational potential of regulating gut microbial metabolic functions as a viable strategy for preventing and managing chemotherapy-induced enterotoxicity. In this study, we demonstrated that DKT repaired intestinal barrier by regulating *Limosilactobacillus-*indole metabolites, and reduced intra-luminal SN-38 production by inhibiting *E. coli* and its β-GUS activity in a rat model of irinotecan-induced enterotoxicity. Notably, these protective effects were achieved without compromising the anti-tumor efficacy of irinotecan, thus highlighting the clinical potential of DKT as a safe adjunctive therapy in chemotherapy settings.

Both preclinical and clinical studies have highlighted the pivotal role of gut microbiota in shaping the enterotoxicity of irinotecan. In our study, *Lactobacillus* and *Limosilactobacillus* constituted nearly 90% of the intestinal microbiota, positioning them as dominant genera that influenced both the progression of enterotoxicity, and the therapeutic response to DKT. Notably, *Limosilactobacillus* was significantly decreased in irinotecan-treated rats but reversed by DKT, whereas *Lactobacillus* remained largely unaffected by DKT. These findings suggest that DKT may protect against enterotoxicity primarily by enriching *Limosilactobacillus*. *L. reuteri* was further identified as the dominant bacterial species contributing to the therapeutic benefit of DKT. However, a recent study showed that *L. reuteri* can exacerbate irinotecan–induced enterotoxicity by producing β-GUS^47^. The inconsistency between the results may be due to the different bacterial strains used in the two studies. *Lactobacillus* was significantly elevated in the model group, which is interesting given that it plays beneficial roles in intestinal homeostasis^48^. Nevertheless, a similar increase was observed in *Lactobacillus* following CPT-11 intervention, which was also positively correlated with Cox-2, IL-6, FITC-dextran, and LPS levels, and negatively correlated with ZO-1 expression^34^. Consistent with this, a metagenomic study revealed that *Lactobacillus* was enriched in the patients receiving CPT-11 treatment^49^. Furthermore, in chemotherapy-induced mucositis, *Lactobacillus* abundance surged from 5.32% in the untreated controls to 26.08% after irinotecan and 5-fluorouracil treatment^50^. Altogether, these findings suggest a context-dependent role of *Lactobacillus* in irinotecan-induced enterotoxicity, possibly distinct from its conventional probiotic functions. Nevertheless, given the phylogenetic and functional diversity within the genus *Lactobacillus*, the precise contributions of individual species to irinotecan-induced enterotoxicity remain poorly understood. Therefore, a comprehensive, species-level characterization of gut microbial dynamics in patients receiving irinotecan is urgently warranted. To investigate the underlying microbiota–dependent mechanisms, KEGG enrichment analysis was performed and identified the endocannabinoid signaling pathway as a key candidate mediating the effects of DKT. Consistent with this, the main receptor CB1 but not CB2 was normalized by DKT at both the transcriptional and protein levels. Given that CB1 is predominantly expressed in the intestinal epithelium, we speculated that intestinal bacteria may protect the barrier through CB1 signaling.

Gut microbial metabolites function both as signaling mediators in the host-microbiota crosstalk and as dynamic markers of disease states. Tryptophan metabolism has emerged as a key factor in driving irinotecan-induced enterotoxicity^9,18,51^. Herein, we found that irinotecan intervention disrupted tryptophan metabolism, as indicated by the downregulation in IPA and IE, and upregulation in IAA, ILA, XA, and other metabolites. This metabolic profile was consistent with previous clinical observations and untargeted metabolomics studies, which revealed notable enrichment of the tryptophan metabolomic pathway. Furthermore, IAA levels are elevated in both feces and serum of diarrheal patients receiving irinotecan, which correlates significantly with the severity of diarrhea^9^. However, clinical data on other tryptophan metabolites remain limited. In our rat model of enterotoxicity, ILA level was elevated by more than three-fold, presumably as a result of the expansion of ILA-producing bacteria such as *Lactobacillus* following CPT-11 treatment^42,52^. ILA is known to exert protective effects in colitis and CRC models by activating AhR signaling, which reinforces the intestinal barrier and attenuates inflammatory responses^53,54^, which may account for the elevated IL-10 levels of enterotoxicity rats in Fig. S1E. However, its role appears highly context-dependent. Specifically, in cisplatin-induced intestinal injury model, increased serum ILA levels were strongly associated with impaired intestinal barrier function^55^. Moreover, exogenous ILA administration exacerbated inflammatory symptoms in colitis model through autophagy inhibition, and reversed the anti-colitis effects of the total glucosides of peony^56^. XA level exhibited similar trends and fold changes to that of ILA following irinotecan intervention. The transformation from ulcerative colitis to colitis-associated CRC is accompanied by a progressive increase in XA level across three pathological stages: inflammation, dysplasia, and carcinoma^57^. In our study, both ILA and XA were positively correlated with *Lactobacillus* and negatively correlated with *Limosilactobacillus,* indicating their potential detrimental role of these metabolites in irinotecan-induced enterotoxicity. Therefore, the functional roles of ILA and XA may depend on the specific pathological context, and the precise biological significance of tryptophan metabolites in irinotecan-induced enterotoxicity needs further investigation. DKT intervention normalized the perturbations in tryptophan metabolism, and the normalization of IPA and IE levels were attributed to the enrichment of *L. reuteri*, which subsequently improved intestinal barrier integrity^42,43^. Consistent with our findings, IPA is positively associated with the abundance of *L. reuteri* (*P* = 0.0083, *r* = 0.3767) and activates pregnane X receptor to protect against 5-fluorouracil-associated diarrhea and intestinal barrier damage^58^. In addition, the AhR agonist IE protected against colitis and regulated intestinal barrier function in colitis model^59,60^. These findings suggest that IPA and IE are key functional metabolites through which DKT exerts its protective effects against irinotecan-induced enterotoxicity.

The pivotal role of gut bacterial β-GUS in irinotecan-induced enterotoxicity is widely acknowledged^12,34^. Furthermore, β–GUS subtypes exhibit marked differences in hydrolytic capacity, with notably higher activity reported for Loop 1 β–GUS derived from *E. coli*, *F. prausnitzii*, and *E. faecalis* within the human colon^34,44,61^. Consistent with these findings, the abundance of Loop 1 β–GUS-producing species significantly increased upon CPT-11 intervention, and was normalized by DKT treatment. This suggests that DKT alleviates enterotoxicity, at least in part, by suppressing the growth of bacteria that produce highly active β–GUS. CPT-11 intervention in specific-pathogen-free mice has been associated with inflammatory responses, epithelial damage, and barrier dysfunction, whereas germ–free animals exhibit only minimal pathological changes in response to CPT-11^17^. In our study, ABX-mediated depletion of the gut microbiota led to mild weight loss and intestinal damage, and no signs of diarrhea were observed. The beneficial effects of the absence of gut bacteria can be largely attributed to the lower level of bacterial β–GUS, which is responsible for converting inactive form SN–38G to active form SN–38. However, while monocolonization of germ-free mice with *E. coli* increased intestinal permeability, it failed to reproduce the full symptoms of irinotecan-induced enterotoxicity, including neutrophilic infiltration or epithelial disruption^17^. This suggests the involvement of additional microbiome-driven mechanisms, such as gut microbiota-mediated tryptophan metabolism, and underscores the need for a more comprehensive investigation of gut microbiota functions.

The entero-protective effect of DKT was achieved via metabolic remodeling of the gut microbiota, encompassing both endogenous tryptophan metabolism and the intestinal metabolism of SN-38G. Multiple DKT components were identified that likely restored intestinal tryptophan metabolism in the irinotecan-induced enterotoxicity model. This is supported by the fact that the herbs in DKT, including Sichuan pepper, dried ginger, and ginseng, can regulate gut microbial tryptophan metabolism in different disease models^62,63^. Specifically, in 5-fluorouracil-induced intestinal mucositis model, the essential oil derived from dried ginger modulates tryptophan–mediated AhR signaling axis to against mucositis^64^. In addition, GRg_1_ from ginseng is known to regulate gut microbial tryptophan metabolism, and improve the intestinal barrier and inflammation in colitis mice^65^. We further evaluated the effects of eight main components of DKT on *E. coli* and its Loop 1 β-GUS, and found that HαS, HβS and 6-SGL significantly inhibited the growth of *E. coli* and selectively suppressed Loop 1 β-GUS activity. However, these components were ineffective against mammalian β-GUS activity, indicating that DKT can inhibit the production of SN-38 without affecting the normal physiological processes. Notably, 6-SGL exhibited a superior inhibitory effect against Loop 1 β-GUS compared to DSA at the same concentration. Apart from its direct effects on microbial metabolism, 6-SGL also possesses anti-inflammatory and antineoplastic characteristics^66,67^. Furthermore, *in vitro* data show that 6-SGL acts synergistically with irinotecan against colon cancer cells^68^. Thus, 6-SGL is a promising compound for mitigating irinotecan-induced enterotoxicity.

Taken together, DKT reversed gut dysbiosis in a rat model of irinotecan-induced enterotoxicity, restored tryptophan metabolism, and specifically inhibited Loop 1 β-GUS activity, which mitigated the pathological effects of irinotecan and repaired the intestinal barrier function. The *Limosilactobacillus-*indole metabolites were identified as the functional basis for intestinal barrier injury, and 6-SGL was the key pharmacological component of DKT that inhibited *E. coli* and β-GUS to reduce intra-luminal SN-38 concentration. Our findings clarify a gut microbiota-metabolic remodeling mechanism through which DKT exerts its protective effects against irinotecan–induced enterotoxicity, offering an effective and sustainable strategy for managing this debilitating complication of chemotherapy.

## 5. Conclusions

DKT significantly alleviated irinotecan-induced enterotoxicity in rats by regulating gut microbial metabolism without affecting the anti-tumor effects of irinotecan. The therapeutic efficacy of DKT was primarily attributed to the enrichment of *Limosilactobacillus*-indole metabolites to repair the intestinal barrier, and the inhibition of *E. coli* and its β-GUS activity to reduce SN-38 concentration. Our study underscores the potential of DKT for managing irinotecan-induced enterotoxicity and elucidates the underlying gut microbiota-driven mechanism.

## ^1^Abbreviation

5-HT: serotonin
6-GGL: 6-gingerol
6-SGL: 6-shogaol
ABX: antibiotic cocktail
AhR: aryl hydrocarbon receptor
β-GUS: β-glucuronidase
CB1: cannabinoid receptor 1
CB2: cannabinoid receptor 2
CPT-11: irinotecan
DKT: Daikenchuto
DSA: D-saccharic acid 1,4-lactone hydrate
DSS: dextran sulfate sodium
GRb_1_: ginsenoside Rb_1_
GRe: ginsenoside Re
GRg_1_: ginsenoside Rg_1_
HαS: hydroxy-α-sanshool
HβS: hydroxy-β-sanshool
IAA: indole-3-acetic acid
IAA-Ala: indole-3-acetyl-alanine
IAA-Asp: indole-3-acetyl-aspartate
ICA: indole-3-carboxaldehyde
IE: indole-3-ethanol
IGA: indole-3-glyoxylicacid
IL: interleukin
ILA: indole-3-lactic acid
IPA: indole-3-propionic acid
IS: indoxylsulfate
KEGG: Kyoto Encyclopedia of Genes and Genomes
KYN: kynurenine
LOP: loperamide
NA: nicotinicacid
NAS: N-acetyl-5-hydroxytryptamine;
PCoA: principal coordinate analysis
PLS-DA: partial least squares discriminant analysis
PNPG: 4-nitrophenyl-β-D-glucopyranoside
SMN: skimmianine
SN-38G: 7-ethyl-10-hydroxy camptothecin glucuronic acid
SN-38: 7-ethyl-10-hydroxy camptothecin
Trp: tryptophan
XA: xanthurenic acid.

## Notes

### Competing Interest Statement

The authors have declared no competing interest.

### Funding

- National Natural Scientific Foundation of China: U24A20796, 82174059
- National Leading Talent Project Funding of China: TP-001-2025
- Outstanding young and middle-aged science and technology talents of Shaanxi Administration of Traditional Chinese Medicine: 2023-ZQNY-004

