## Supporting information for "Daikenchuto modulates gut microbial metabolism to mitigate irinotecan-induced enterotoxicity"

### 1. Methods

#### *1.1. Reagents*

Indole-3-lactic acid (ILA), indole-3-acetyl-alanine (IAA-Ala), indole-3-acetyl-aspartate (IAA-Asp), kynurenine (KYN), xanthurenic acid (XA) and serotonin (5-HT) were purchased from Aladdin Biochemical Technology Co., Ltd. (Shanghai, China).

#### *1.2. Immunohistochemistry analysis*

Paraffin slices of small intestine tissue were deparaffinized and rehydrated. The slices were then placed in 3% hydrogen peroxide solution to block endogenous peroxidase and subsequently in 3% bovine serum albumin to closure. The samples were incubated with the primary antibodies overnight at 4°C, followed by co-staining with corresponding secondary antibodies (Cy3 conjugated) for 50 mins at room temperature. The slices were stained with diaminobenzidine chromogenic reagent (Servicebio, Wuhan, China) and were counterstained with hematoxylin for cell nuclei staining.

#### *1.3. Effects of tryptophan metabolites on Loop 1 β-GUS activity*

DSA, ILA, IPA, IE, IAA-Ala, IAA-Asp, XA and 5-HT were prepared as working solution with a concentration of 1 mM using dimethyl sulfoxide solution (DMSO). EcGUS was prepared as working solution with a concentration of 37.5 U/mL using PBS. The final incubation system included 10 uL EcGUS (final concentration: 3.75 U/mL), 10 uL Trp metabolites/DSA/DMSO (final concentration: 100 μM), and 10 uL PNPG (final concentration: 250 μM) and 70 uL PBS. After a 40-minute incubation at 37°C, 70 μL of NaOH solution was added to terminate the reaction, and the absorbance was measured at 405 nm. DMSO was used as a blank control in this experiment and all experiments were repeated for three times.

#### *1.4. Statistical analysis*

Statistical analysis was performed using GraphPad Prism 8.0 (GraphPad, San Diego, CA). Data are presented as the mean ± SEM. Statistical analysis for multiple groups was performed by one-way of variance (ANOVA) followed by Dunnett's *post hoc* multiple comparison's test. The differences were significant at $P < 0.05$.

**Supporting tables**

**Table S1** Sequences of the gene-specific primers

| Gene | Species | Forward primer (5′–3′) | Reverse primer (5′–3′) |
|---|---|---|---|
| *Tjp1* | Rat | AAGATGGGATTCTTAGGCCCAGCA | TCTTTGGCTGCAGGGCTATCTTCT |
| *Ocln* | Rat | ATGTATGGCGGAGAGATGCACGTT | ATAGGCTCTGTCCCAAGCAAGTGT |
| *Cldn1* | Rat | TCTGGGTTTCATCCTGGCTT | TCAGATTCAGCAAGGAGTCGA |
| *Cldn4* | Rat | GGAAGGGTGGAGGTGGTTTA | GGGTAGGGAATTCAGAGGGG |
| *Cnr1* | Rat | AGTGTGCTGCTGCTGTTCATTG | CGTGTGGATGATGATGCTCTTCTG |
| *Cnr2* | Rat | ACCGATACCTATGTCTGTGCTACC | GGAGATCAACGCCGAGAGGAC |
| *Gapdh* | Rat | GGCACAGTCAAGGCTGAGAATG | ATGGTGGTGAAGACGCCAGTA |
| *TJP1* | Human | GGGACAACAGCATCCTTCCA | AGTGTGGTAAGCGCAGCTC |
| *OCLN* | Human | GCCTGGATGACATGGCTGAT | CTCCCTGGCACCGTTGG |
| *CLDN1* | Human | CGATGAGGTGCAGAAGATGA | CCAGTGAAGAGAGCCTGACC |
| *CNR1* | Human | TGGAACTGCGAGAAACTGCA | ACAGAAGCAGTACGCTGGTG |
| *TNF* | Human | ACTCCCAGGTCCTCTTCAAG | TGATGGCAGAGAGGAGGTTG |
| *IL1B* | Human | GCTTGGTGATGTCTGGTCCA | AACACGCAGGACAGGTACAG |
| *IL6* | Human | AGACAGCCACTCACCTCTTC | AGTGCCTCTTTGCTGCTTTC |
| *GAPDH* | Human | TGCACCACCAACTGCTTAG | AGAGGCAGGGATGATGTTC |
| *Tjp1* | Mouse | TTTTTGACAGGGGGAGTGG | TGCTGCAGAGGTCAAAGTTCAAG |
| *Ocln* | Mouse | ATGTCCGGCCGATGCTCTC | TTTGGCTGCTCTTGGGTCTGTAT |
| *Cldn1* | Mouse | GGCTTCTCTGGGATGGATCG | TTTGCGAAACGCAGGACATC |
| *Cldn4* | Mouse | CCACTCTGTCCACATTGCCT | CTTTGCACAGTCCGGGTTTG |
| *Gapdh* | Mouse | ATGGTGAAGGTCGGTGTGAA | TGGAAGATGGTGATGGGCTT |

**Table S2** The parameter information for UPLC-ESI-MS/MS

| Parameter | Value |
|---|---|
| Curtain gas | 30 psi |
| Collision-activated dissociation | medium |
| Ion spray voltage | 5500 V in positive mode |
| | 4500 V in negative mode |
| Source temperature | 450°C |
| Column temperature | 40°C |
| Spray gas | 50 psi |
| Auxiliary gas | 50 psi |

**Supporting figures:**

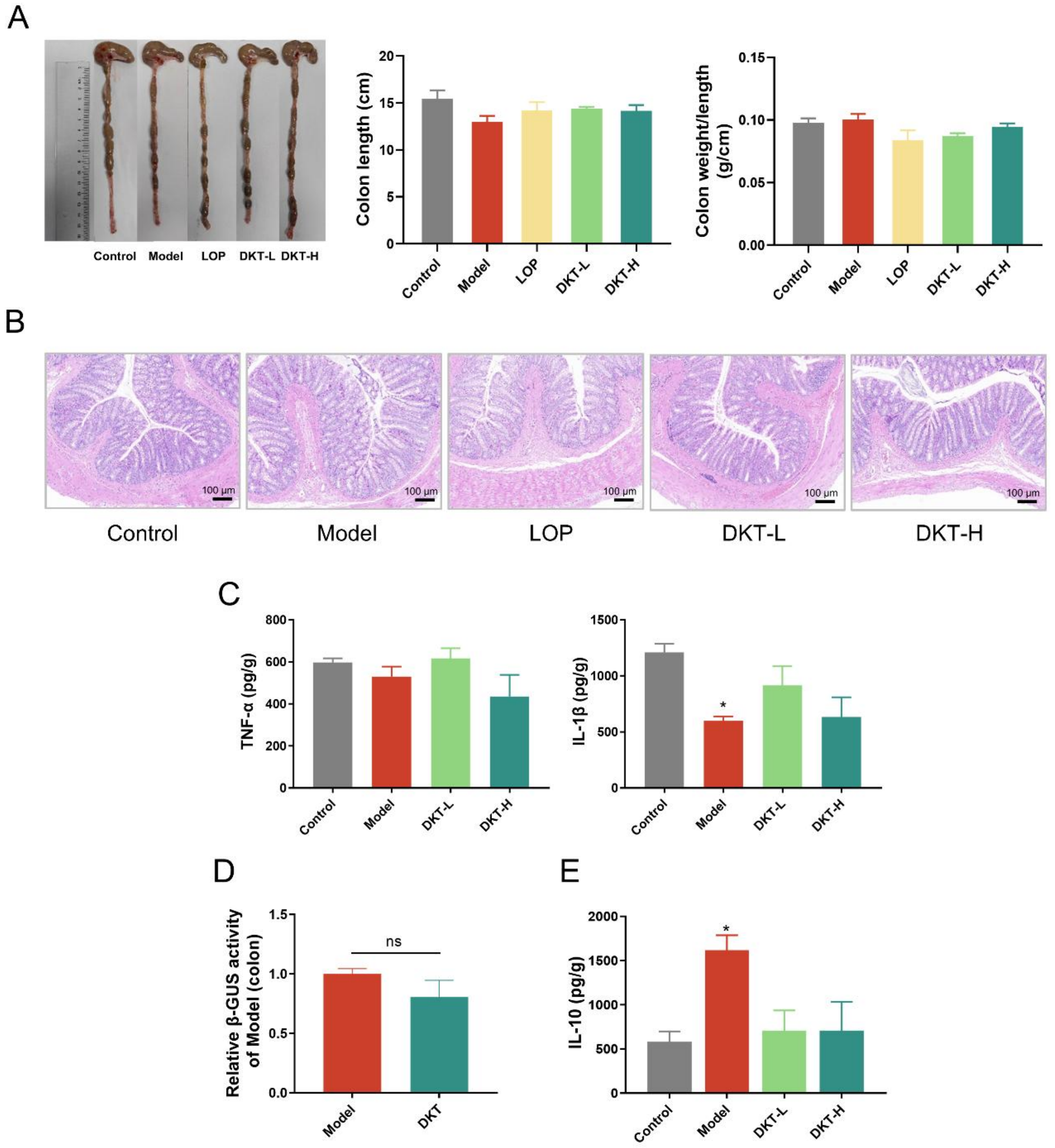


**Figure S1** Evaluation of toxicity in colon tissue of different groups. (A) Representative pictures of colon, colon length and colon weight to length ratio in rats (*n*=6). (B) Representative hematoxylin and eosin (H&E) stained pictures of the colon (scale bars = 100 μm) (*n*=3). (C) Inflammatory cytokine levels (TNF-α and IL-1β) in colon tissues (*n*=3). (D) β-GUS activity in colonic contents of Model and DKT groups (*n*=4). (E) Inflammatory cytokine levels (IL-10) in small intestine tissues (*n*=3). Data are presented as the mean ± SEM. $^{*}P < 0.05$ *vs*. Control group. ns, not significant.

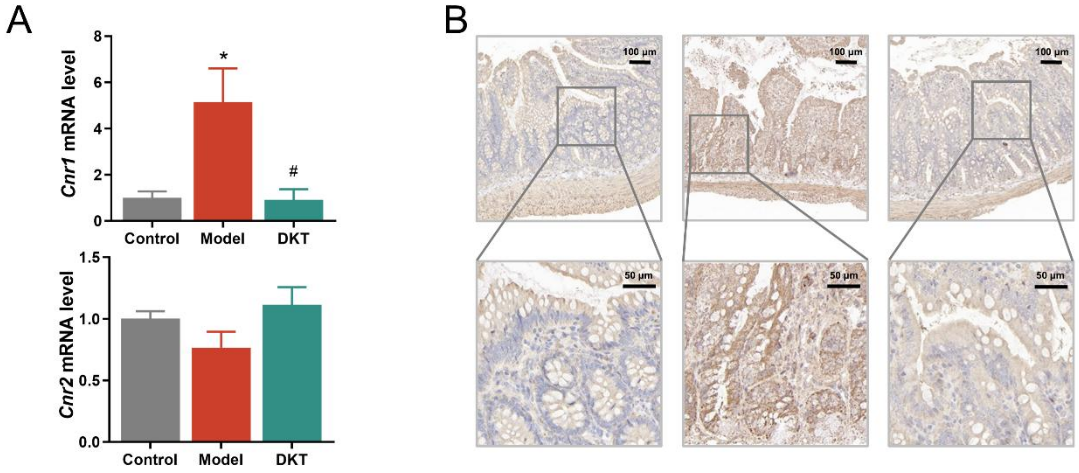


**Figure S2** The effect of DKT on cannabinoid receptors. (A) The mRNA level of *Cnr1* and *Cnr2* in small intestine (*n*=3). (B) Representative images of immunohistochemistry analysis of CB1 in small intestine. Data are presented as the mean ± SEM. $^{*}P < 0.05$ *vs*. Control group, $^{\#}P < 0.05$ *vs*. Model group.

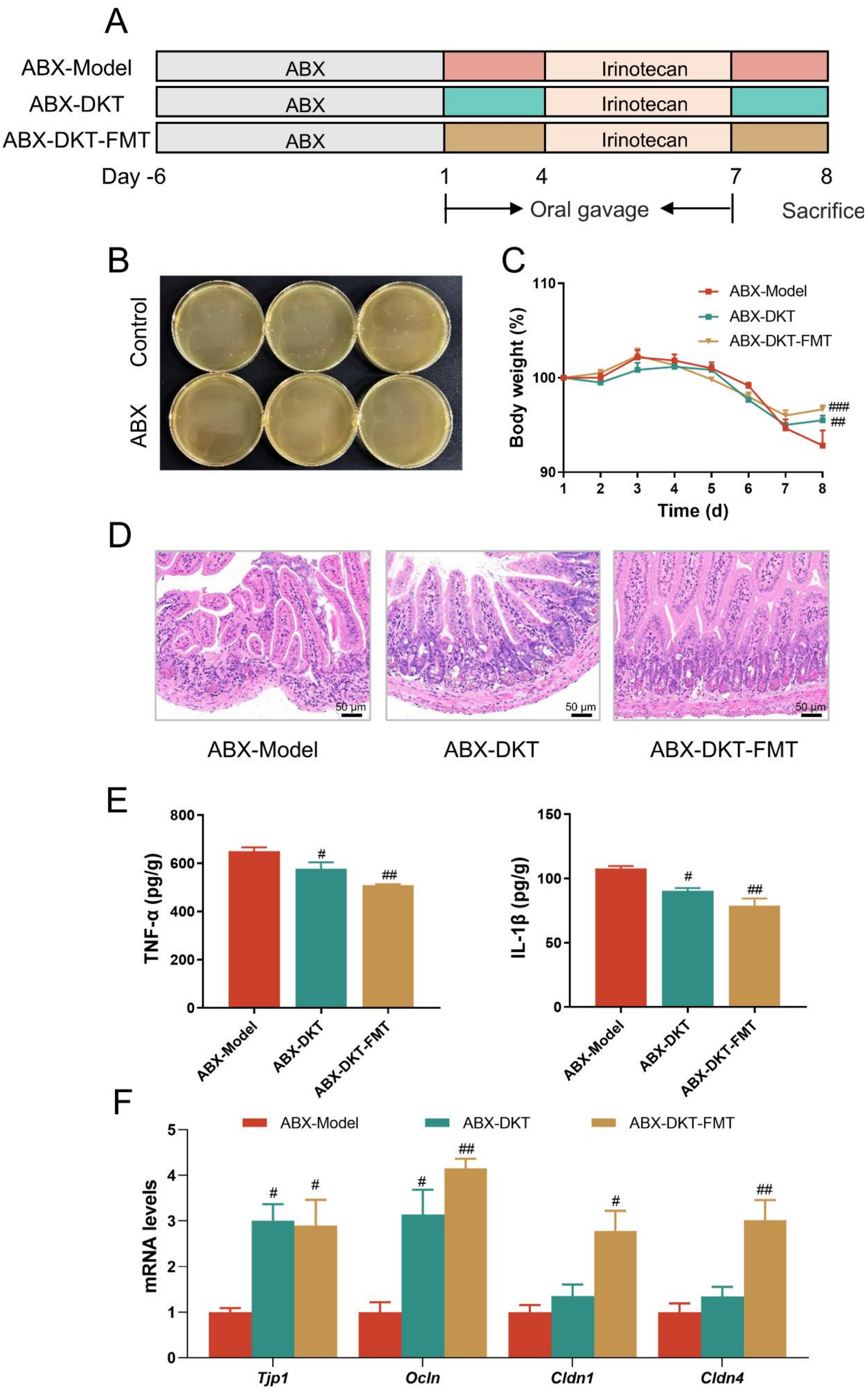


**Figure S3** DKT alleviated the enterotoxicity of irinotecan in a partially dependent manner on gut microbiota

(A) Experimental scheme. (B) Flat coating of fecal microbiota suspension derived from normal and ABX-treated mice. (C) Percentage of body weight during the experiment (*n*=6). (D) Representative hematoxylin and eosin (H&E) stained pictures of the small intestine (scale bars = 100 μm). (E) Inflammatory cytokine levels (TNF-α and IL-1β) in small intestine tissues (*n*=3). (F) The mRNA levels of tight junctions of small intestine tissues (*n*=3). Data are presented as the mean ± SEM. $^{\#}P < 0.05$, $^{\#\#}P < 0.01$ and $^{\#\#\#}P < 0.001$ *vs*. ABX-Model group.

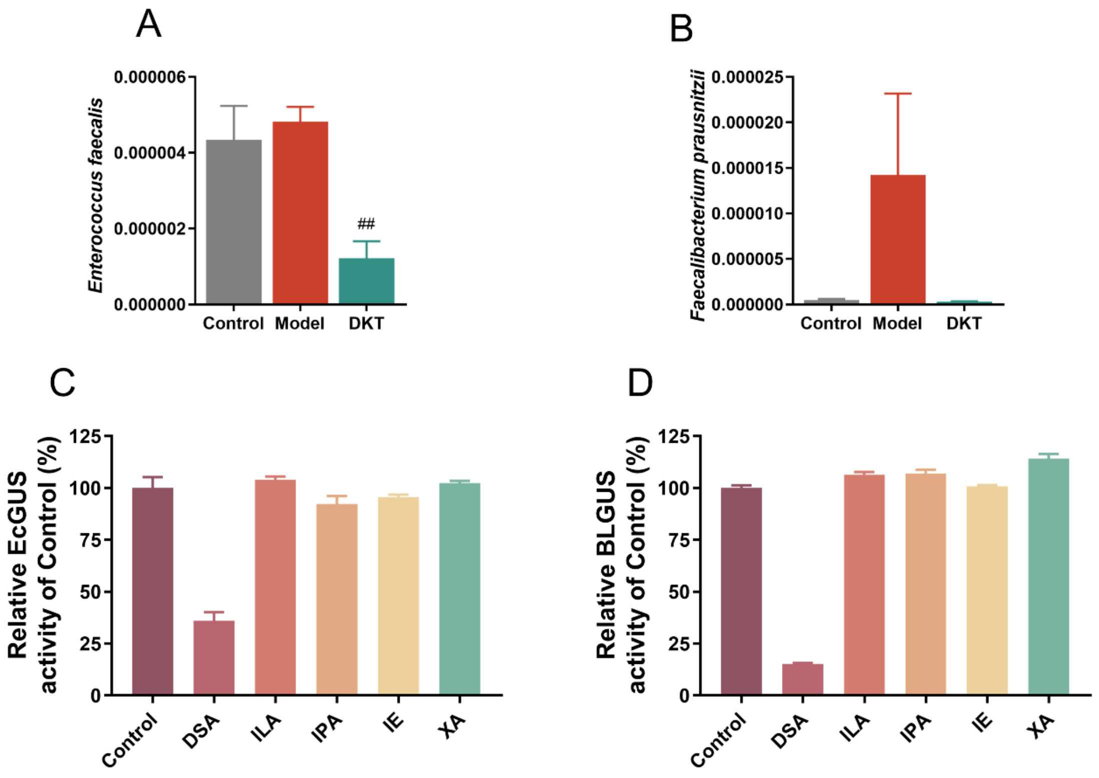


**Figure S4** The relative abundance of *Enterococcus faecalis* (A) and *Faecalibacterium prausnitzii* (B) (*n*=4). Effect of tryptophan metabolites on EcGUS (C) and BLGUS (D) activity (*n*=3). Data are presented as the mean ± SEM. $^{\#\#}P < 0.01$ *vs*. Model group.
